# Actin chromobody imaging reveals sub-organellar actin dynamics

**DOI:** 10.1101/639278

**Authors:** Cara R. Schiavon, Tong Zhang, Bing Zhao, Leonardo Andrade, Melissa Wu, Tsung-Chang Sung, Yelena Dayn, Jasmine W. Feng, Omar A. Quintero, Robert Grosse, Uri Manor

## Abstract

The actin cytoskeleton plays multiple critical roles in cells, from cell migration to organelle dynamics. The small and transient actin structures regulating organelle dynamics are difficult to detect with fluorescence microscopy. We developed an approach using fluorescent protein-tagged actin nanobodies targeted to organelle membranes to enable live cell imaging of previously undetected sub-organellar actin dynamics with high spatiotemporal resolution. These probes reveal that ER-associated actin drives fission of multiple organelles including mitochondria, endosomes, lysosomes, peroxisomes, and the Golgi.

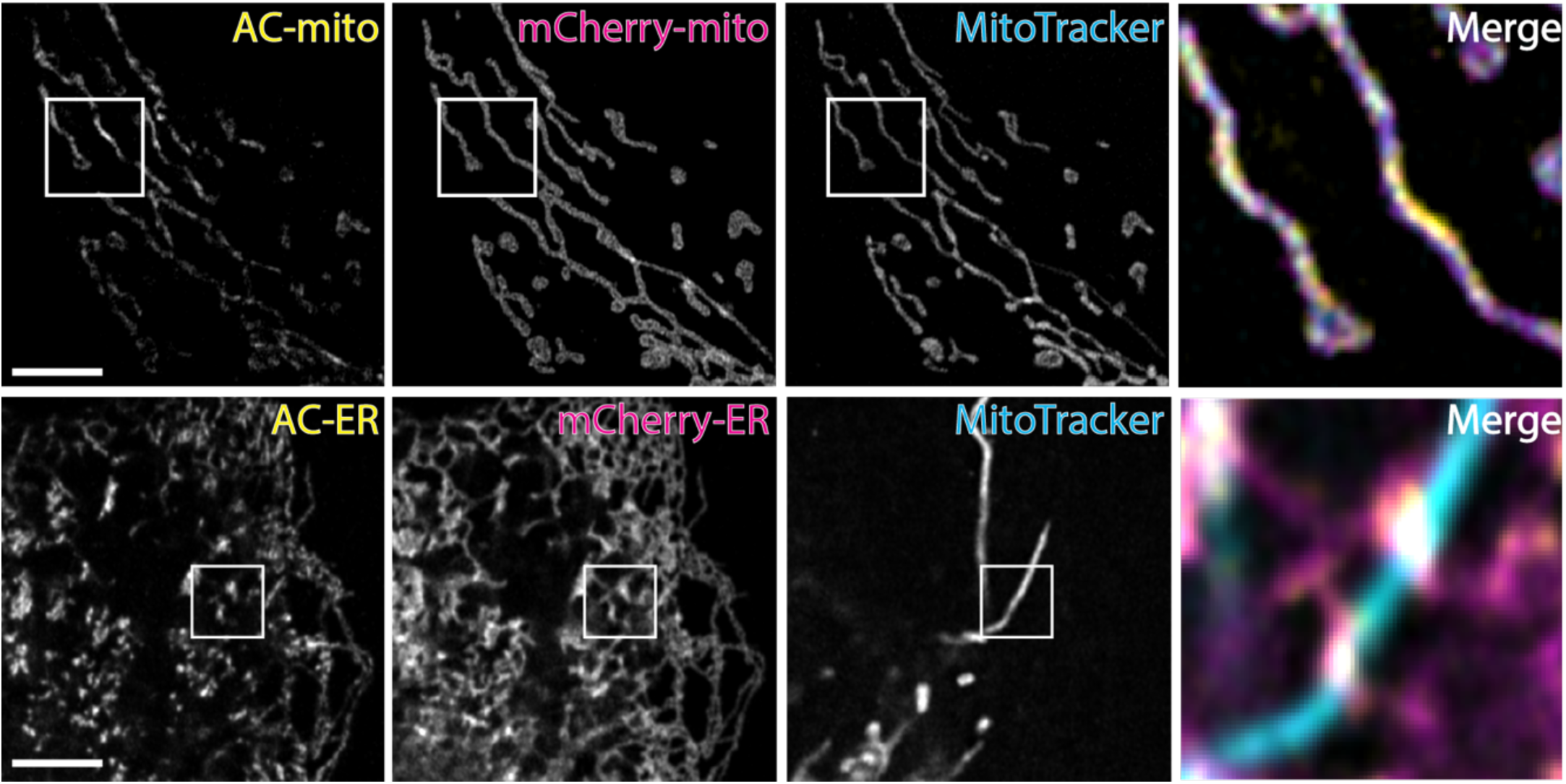

## Introduction

The critical role of the actin cytoskeleton in organelle dynamics is largely accepted, but poorly understood. The precise spatiotemporal dynamics of actin at organelle membranes remain particularly unclear due to the combined limitations of currently available actin probes and imaging approaches. For fluorescence microscopy approaches, imaging smaller actin structures in the cell suffers from an enormous background signal issue - the high signal from the dense meshwork of actin filaments throughout the cell overwhelms the signal from the relatively small, transient actin structures associated with organelle dynamics. Furthermore, the limitations in resolution make it difficult to determine whether any actin filaments are physically associated with the organelle. Meanwhile, EM approaches can be used to visualize actin filaments with extremely high resolution^1,2^. However, these techniques only capture single timepoints, making it difficult to determine the precise state or dynamics of the actin filaments or their associated organelles at the time of fixation. All these limitations ultimately preclude a solid understanding of the mechanisms by which actin regulates organelle dynamics in health or disease. Here we employ fluorescent protein-tagged actin nanobodies, aka “actin chromobodies” (AC)^3,4^, fused to organelle membrane targeting sequences to facilitate live cell imaging of sub-organellar actin dynamics with high spatiotemporal resolution. Using these probes we discovered that ER-associated actin accumulates at all ER-organelle contacts during organelle fission.

## Results

We hypothesized that AC probes with organelle membrane targeting sequences could be used to monitor actin dynamics exclusively within a ∼10nm distance from the target organelle membrane. Given the high mobility of membrane anchored proteins in a bilayer, an F-actin binding probe containing only a membrane targeting sequence and a fluorescent protein tag should quickly accumulate at sites of F-actin enrichment near the membrane (Figure 1a). To test this hypothesis, we generated constructs containing the yeast Fis1 mitochondrial outer membrane and Cytb5ER endoplasmic reticulum (ER) minimal C-terminus tail membrane targeting sequences fused to the cytoplasm-facing actin nanobody and tagGFP (“AC-mito” and “AC-ER”)^5^. Live cell Airyscan imaging of cells expressing AC-mito counterstained with MitoTracker dye revealed strikingly specific regions of AC-mito enrichment on the surfaces of mitochondria (Figure 1b and Supplementary Movies 1-2). Similarly, AC-ER expressing cells revealed significant accumulation of AC-ER at specific ER-mitochondria contact sites (Figure 1b and Supplementary Movies 3-4). To rule out the possibility that the membrane targeting sequences we used were causing the probe to accumulate in specific regions independent of actin-binding activity, we co-transfected AC-mito or AC-ER with mCherry-tagged versions of the mitochondrial (Fis1) and ER (Cytb5ER) membrane targeting sequences (mCherry-mito and mCherry-ER). As expected, the mCherry-mito and mCherry-ER signals were evenly distributed along their respective organelle membranes, with no obvious enrichment in any specific regions. In contrast, the co-expressed AC-mito and AC-ER constructs were uniquely enriched in specific regions on their respective organelles (Figure 1b). mCherry-ER showed some accumulation at ER-mitochondria intersections, indicative of stable contacts at these sites^6^. However, the AC-ER probe showed particularly enhanced enrichment at ER-mitochondria contacts, revealing significant accumulation of actin specifically at these contact sites (Figure 1b). Fluorescence recovery after photobleaching (FRAP) experiments showed that both the AC and mCherry probes are highly mobile on the membrane, but the AC probes exhibit significantly lower mobility and a higher immobile fraction (Figure 1c, Supplementary Movies 5-6), consistent with the model in which a fraction of AC probes are immobilized by actin (Figure 1a).

**Figure 1.**
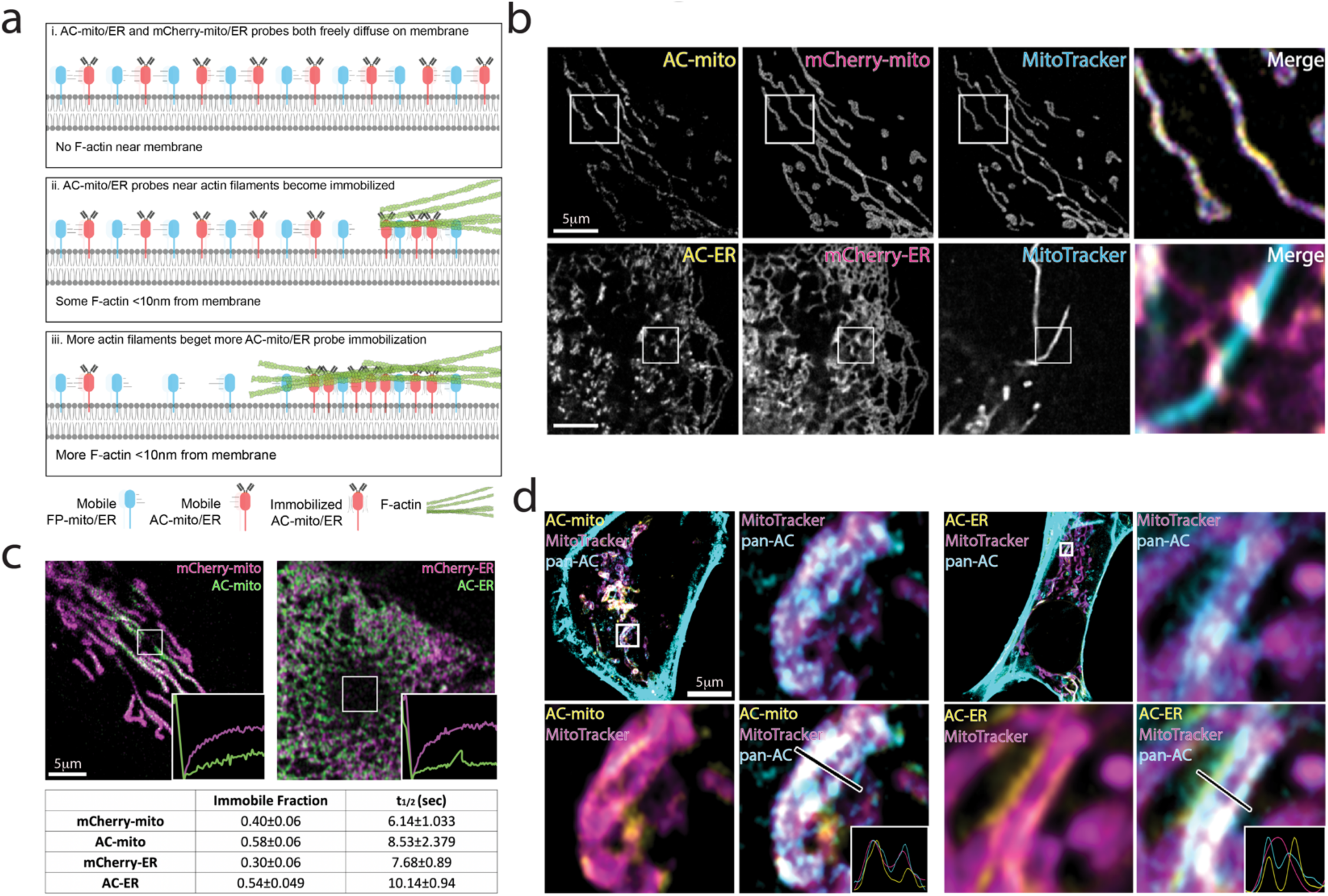
Mitochondria and ER localized actin nanobodies reveal sub-organellar accumulation of actin filaments. **a**,Cartoon depicting change in AC—mito/ER localization in response to the presence of actin. When there is no actin near the membrane, AC-mito/ER freely diffuse throughout the membrane. When actin comes within 10nm of the membrane, only the AC probes are immobilized at the sites of actin accumulation. As actin filaments grow/accumulate, more of the AC probe becomes immobilized. **b**, Co-expression of mCherry-mito/ER in U2OS cells demonstrates AC-mito/ER sub-accumulation depends on actin nanobody activity. Sub-accumulation was observable in all cells imaged (AC-mito: n = 68, AC-ER: n = 55). Scale bars: 5 μm. **c**, FRAP of cells co-expressing AC/mCherry-mito/ER shows that both probes are mobile, but the AC probes have lower mobility and a higher immobile fraction. **d**, Co-expression of a pan-actin label (ACtagRFP) with AC-mito/ER reveals actin is enriched at sites labeled by AC-mito/ER probes.

In our hands, pan-actin probes such as phalloidin fail to clearly reveal sites of actin accumulation on mitochondria or the ER under normal imaging conditions (Supplementary Figure 1). Furthermore, given the abundance of actin signal throughout the cell and the limitations of resolution in fluorescence microscopy, it is extremely difficult to assess whether colocalization of a pan-actin signal and an organelle membrane is a true actin-organelle association or not. Since our AC probes are small proteins tethered to their respective organelle membranes, we can conclude that all observed actin accumulation is within a maximum distance of 10nm from the organelle membrane (although the distal ends of associated actin filaments may extend far beyond 10nm). Using our AC probes as a guide, we were able to identify regions of actin accumulation using a cytoplasmic pan-actin probe (i.e. AC-tagRFP) (Figure 1d). Since phalloidin is widely considered the gold standard for F-actin labeling^4^, we attempted to colocalize AC-mito and AC-ER with phalloidin staining in fixed cells (Supplementary Figure 2). Interestingly, we found that fixation with 4% PFA causes a dramatic loss in AC-mito enrichment as well as an overall decrease in AC-mito or AC-ER signal, perhaps reflective of the relatively small and transient nature of these membrane-associated actin structures. However, we were still able to detect co-accumulation of our probes with phalloidin in fixed cells. Notably, the contrast in the phalloidin signal needed to be extremely high in order to visualize the signal at AC enriched sites (Supplementary Figure 2), causing neighboring structures to be completely saturated or clipped. We next tested whether other fixation methods could both improve preservation of the AC probe signal as well as phalloidin labeling of AC accumulation sites. Inspired by recent work in the Svitkina lab^1^, we found that using a cytoskeletal stabilization buffer and light saponin-mediated membrane extraction better preserved the AC signal and improved phalloidin labeling at AC-mito/ER enrichment sites (Supplementary Figure 2). These results highlight the value of live imaging labeling approaches and show why similar actin enriched sites were not previously detected with standard phalloidin staining and fluorescence imaging. Indeed, only with the AC-mito/ER signal serving as a guide could we confidently identify sub-organellar regions of actin enrichment in our phalloidin signal.

We considered it important to determine whether the Fis1 membrane targeting sequence in our AC-mito probe might induce dominant negative effects on mitochondrial dynamics, perhaps by outcompeting endogenous Fis1 for mitochondrial outer membrane localization. Since the AC-mito construct uses only the C-terminal membrane targeting domain of Fis1, an anti-Fis1 antibody against the cytoplasmic N-terminal domain should only detect endogenous protein. We found no change in endogenous Fis1 localization to mitochondria in AC-mito expressing cells compared to neighboring non-transfected cells (Supplementary Figure 3). Finally, to determine whether our localization results were specific to the Fis1 membrane targeting sequence or to actin nanobodies in particular, we generated variants of our AC-mito probe using a different membrane targeting sequence (Cytb5mito^5^) and/or a different F-actin probe (LifeAct^7^). All four of our probes yielded very similar results, revealing specific sub-mitochondrial regions of actin accumulation (Supplementary Figure 4). However, the LifeAct probes displayed a more diffuse pattern on the mitochondrial outer membrane. In contrast, our AC probe seemed to better highlight specific subdomains of actin enrichment, consistent with previous comparative studies suggesting that actin nanobodies have superior F-actin labeling capabilities^4^. Thus, we decided to mainly use our AC probes for the majority of our experiments. To further test whether AC-mito accumulation on mitochondria is dependent on F-actin, we treated AC-mito expressing cells with the F-actin depolymerizing drug Latrunculin B (LatB), which significantly reduced the AC-mito signal. Most of the remaining signal appeared as punctate structures, matching the punctate F-actin structures observed with the pan-actin probe AC-tagRFP (“pan-AC”) (Supplementary Figure 5). Importantly, LatB treatment resulted in a more diffuse distribution of AC-mito, reflective of a reduction of F-actin available on or near mitochondria to induce accumulation. Similar results were observed after treatment with cytochalasin D (Supplementary Movie 7). Taken together, these results strongly support the conclusion that our AC probes are labeling F-actin on their respective membrane surfaces. These probes thus facilitate unprecedented resolution (10nm from the membrane) and signal-to-background imaging of mitochondria- and ER-associated actin filaments in live cells. It should be noted that most (but not all – see Moore et al.^8^) previous live imaging of actin and mitochondria was performed during overexpression or stress conditions in order to increase mitochondrial fission and/or the associated actin signal at mitochondria^9-14^, whereas these data were obtained under normal physiological conditions.

Previous studies have demonstrated a role for ER-mitochondria contact sites in driving mitochondrial fission, likely with actin playing a role in force generation and Drp1 recruitment and activation^1,9-15^. However, due to the abundance of actin in the cytoplasm, it is difficult to detect actin accumulating near ER-mitochondria contact sites, or whether any such actin directly associates with either organelle. Furthermore, pharmacological or genetic perturbation of actin dynamics likely causes global changes to the entire cell, making it difficult to distinguish direct versus indirect effects. To determine whether mitochondria- or ER-associated actin accumulates during ER-mediated fission, we performed live imaging of cells co-expressing AC-mito and mCherry-tagged Drp1, an actin-binding dynamin-related GTPase protein essential for mitochondrial fission^10,16,17^ (Figure 2, Supplementary Figure 6, and Supplementary Movies 8-11). To avoid potential overexpression artifacts, we used the low expression ubiquitin promoter C to drive expression of our AC probes^18^. We found AC-mito accumulates at or immediately adjacent to mitochondrial fission sites prior to Drp1-mediated fission (Figure 2a, Supplementary Figure 6a, Supplementary Movie 8). Interestingly, we frequently observed elongated regions of AC-mito enrichment crossing over mitochondrial constriction sites, consistent with recent reports of similar actin structures observed with platinum replica electron microscopy of membrane extracted cells^1^. We similarly observed accumulation of AC-ER at mitochondrial fission sites prior to Drp1-mediated fission (Figure 2b, Supplementary Figure 6b, Supplementary Movie 9). Co-expression of AC-mito or AC-ER with mCherry-ER showed that both mitochondria- and ER-associated actin accumulates at ER-tubules marking mitochondrial fission sites (Figure 2c-d, Supplementary Figure 6c-d, Supplementary Movies 10-11). Finally, co-expression of AC-mito and AC-ER revealed a consistent pattern wherein mitochondrial-associated actin accumulates first, followed by ER-associated actin prior to fission (Figure 2e, Supplementary Figure 6e, Supplementary Movie 12). These results clarify the spatiotemporal dynamics of a subpopulation of actin filaments specifically associated with the mitochondrial outer membrane versus the ER prior to and during mitochondrial fission.

**Figure 2.**
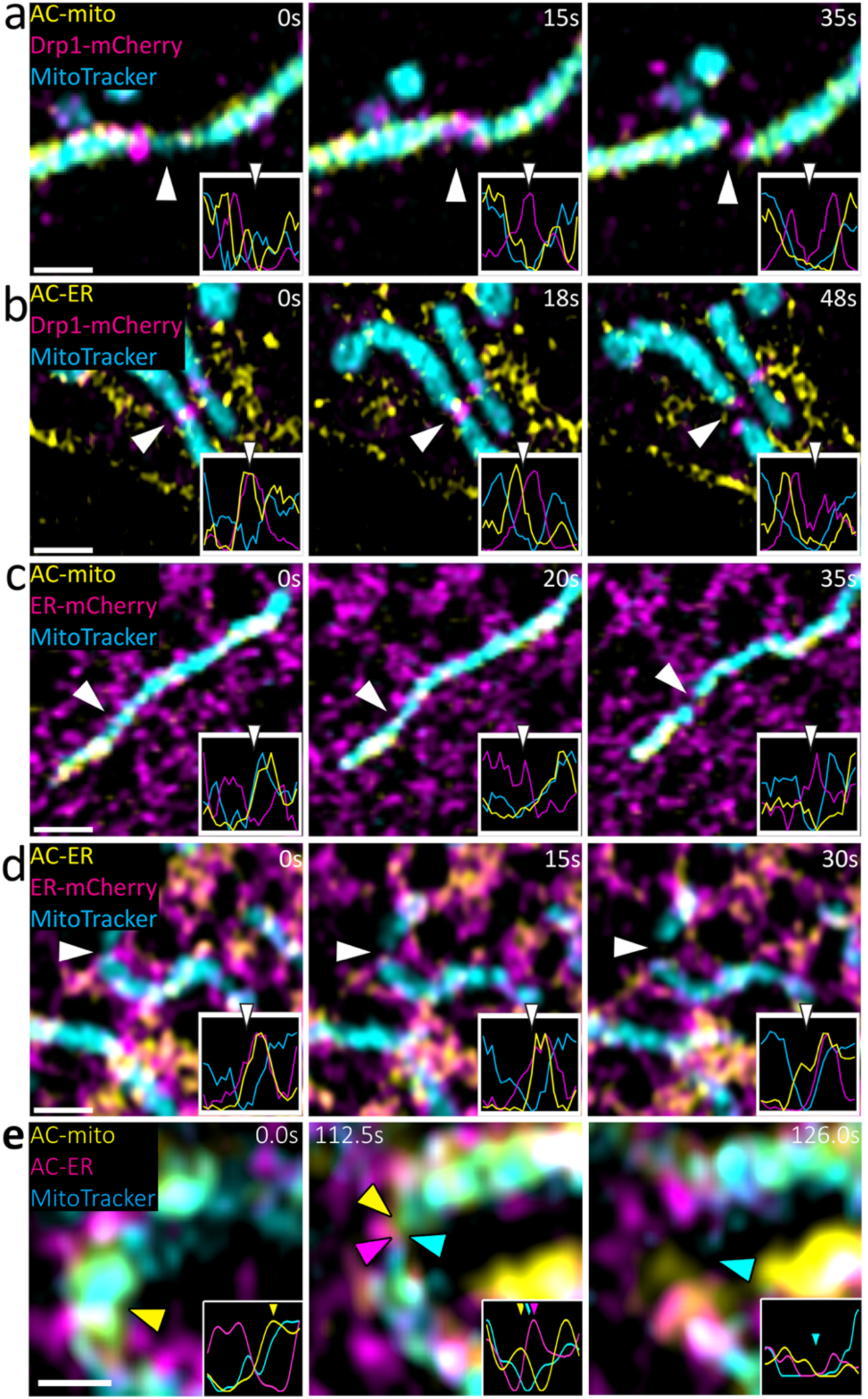
Mitochondria- and ER-associated actin accumulates during mitochondria fission. **a,** AC-mito accumulates at mitochondrial fission sites prior to Drp1. **b,** AC-ER co-accumulates with Drp1 during mitochondrial fission. **c,** AC-mito accumulates at ER-mediated mitochondrial fission sites. **d,** AC-ER accumulates at ER-tubules driving mitochondrial fission. Scale bars: 1 μm. **e,** AC-mito accumulates at mitochondrial fission sites prior to AC-ER accumulation. Scale bar: 0.5 μm. In total, AC-mito and AC-ER accumulation was observed at nearly all mitochondrial fission events (AC-mito: 32/33 fission events, AC-ER: 31/31 fission events).

Recent studies have suggested an analogous role for the ER in mediating endosomal fission^19,20^. However, direct evidence for ER-associated actin during fission is missing. Using our AC-ER probe, we observed actin enrichment at ER-associated actin at endosomal constriction and fission sites (Figure 3b, Supplementary Movie 14). Given the ER regularly contacts most organelles in the cell more than 90% of the time^21^, we hypothesized that actin accumulates at all ER-organelle contact sites to drive their constriction as a preparatory step for fission. Live imaging of cells co-expressing AC-ER and markers for mitochondria, endosomes, peroxisomes, lysosomes, and the Golgi all revealed accumulation of ER-associated actin at fission sites for all of these organelles (Figure 3a-e, Supplementary Movies 13-17). Importantly, the frequency of AC-ER probe accumulation was significantly greater than what would be expected by chance (Supplementary Figure 7). Scanning electron microscopy (SEM) imaging of cells treated with saponin and cytoskeleton stabilization buffer shows actin filaments associated with mitochondria in similar patterns observed with AC-mito (Figure 3f). Finally, AC-ER reveals a striking network of ER-associated actin bundles on the nucleus (Figure 3g, Supplementary Movie 18). SEM imaging of non-transfected cells treated with saponin and cytoskeleton stabilization buffer shows a similar network of actin fibers accumulating on the nucleus, validating the localization pattern observed in our AC-ER fluorescence data (Figure 3g). A perinuclear actin cap structure has been previously reported and is known to have important roles in multiple cellular processes^22-27^. However, as with other actin structures, the actin filaments specifically associated with the nucleus have been difficult to visualize with pan-actin probes due to the overwhelming signal from surrounding actin stress fibers and cortical actin. Thus, AC-ER labeling provides a more specific labelling approach for visualization of perinuclear actin structures.

**Figure 3.**
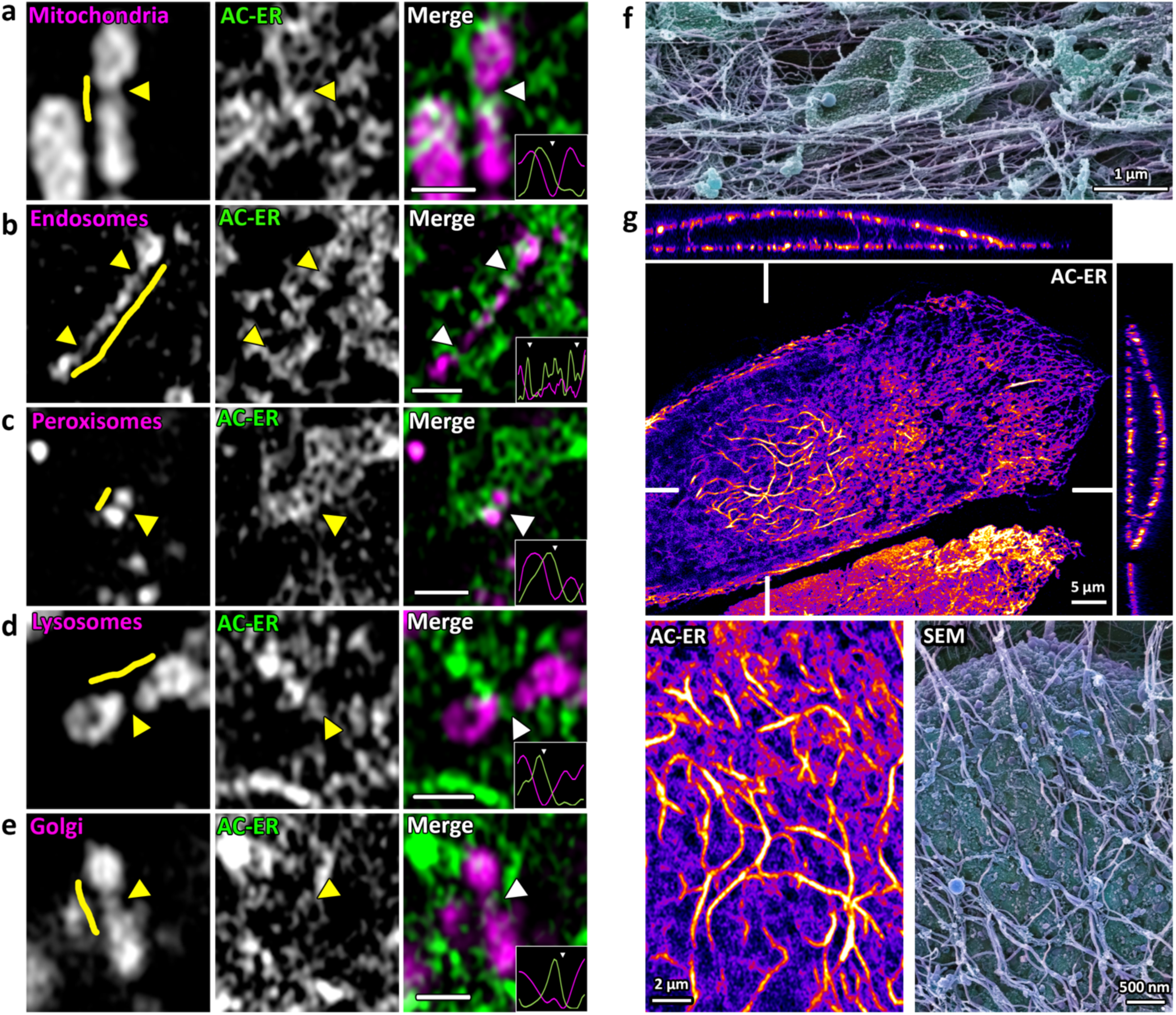
ER-associated actin accumulates at most organelle fission sites. AC-ER accumulates at fission sites in (**a)** mitochondria; **(b)** endosomes (labeled with Rab5a-mCherry); (**c)** peroxisomes (labeled with dsRed-Skl); **(d)** lysosomes (labeled with LAMP1-mCherry); and **(e)** Golgi (labeled with SiT-mApple). AC-ER accumulation was observable at nearly all fission/constriction events imaged (mitochondria: 31/31, endosomes: 17/17, peroxisomes: 12/12, lysosomes: 21/22, Golgi: 6/6. **(f)** SEM image of actin filament association with a mitochondrion. **(g)** AC-ER reveals a distinct pattern of ER-associated actin accumulation around the nucleus. This pattern was observed in 16/22 cells expressing AC-ER. SEM imaging of membrane extracted cells reveals similar structures. Scale bars for a-e: 1 μm

## Conclusion

Taken together, our data reveal a novel role for the ER in recruiting actin filaments to all ER-organelle membrane contact sites, most likely in order to drive membrane remodeling and scaffolding for dynamin-mediated membrane scission. Given the important role of both organelle-organelle contact sites and the actin cytoskeleton in health and disease, the apparent ability of the ER to drive actin accumulation at organelle contacts has broad implications for both cell biology and biomedical research. Finally, the ability to use membrane-anchored AC probes as “proximity sensors” for sub-cellular actin dynamics provides a novel tool for studying the role of actin in a wide range of cell biological processes. The variety of subcellular compartments, protein targets, and corresponding subcellular or sub-organellar processes that can be studied will increase as additional nanobodies and targeting sequences are developed.

## Methods

### Cell culture

U2OS and Hap1 cells were purchased from ATCC. Cells were grown in DMEM supplemented with 10% fetal bovine serum at 37°C with 5% CO_2_. Cells were transfected with Lipofectamine 2000 (ThermoFisher). Cells were plated onto either 8-well #1.5 imaging chambers or #1.5 35mm dishes (Cellvis) that were coated with 10μg/mL fibronectin in PBS at 37°C for 30 minutes prior to plating. 50nM MitoTracker Deep Red (ThermoFisher) was added for 30 minutes then washed for at least 30 minutes to allow for recovery time before imaging in FluoroBrite (ThermoFisher) media.

### Airyscan confocal imaging

Cells were imaged with a 63× 1.4NA oil objective on a ZEISS 880 LSM Airyscan confocal system with an inverted stage and heated incubation system with 5% CO_2_ control. The GFP channels were imaged with a 488nm laser line at ∼500nW laser power. The mCherry or tagRFP channels were imaged with 561nm laser at ∼1μW laser power. The MitoTracker Deep Red channel was imaged with ∼250nW laser power. For timelapse imaging, the zoom factor was set between 3x-6x to increase the frame rate. In all cases, the maximum pixel dwell time (∼0.684μs/pixel) and 2x Nyquist optimal pixel size (∼40nm/pixel) was used.

### Cytoskeleton Stabilized Membrane Extraction of Cells for Fluorescence Imaging

Cells were cultured and imaged as described above. Medium was replaced with cytoskeleton stabilization membrane extraction buffer (0.05% saponin, 100mM PIPES, pH 6.8, 1mM EGTA, 1mM MgCl_2_, 4.2% sucrose, 1mM PMSF, 10μM taxol, 10μM phalloidin-Alexa568) pre-warmed to 37°C. Cells were imaged immediately before and 10min after the addition of membrane extraction buffer.

### Antibodies

We used the rabbit anti-Fis1 antibody against the N-terminus cytoplasmic facing side of the human Fis1 protein, made by Prestige Antibodies Powered by Atlas Antibodies (Sigma-Aldrich, catalog #: HPA017430). The amino acid sequence of the antigen is MEAVLNELVSVEDLLKFEKKFQSEKAAGSVSKSTQFEYAWCLVRSKYNDDIRKGIVLLEELLPKGS KEEQRDYVFYLAVGNYRLKEYEKALKYVRGLLQTEPQNNQAKELERLIDKAMKKD.

### Immunofluorescence

Cells were washed in PBS then fixed with 4% PFA for 30 minutes before permeabilization with 0.1% Triton X-100 for 30 minutes. Cells were then blocked overnight with 4% BSA at 4°C. Cells were then incubated with primary antibody for 2 hours, rinsed 3x with PBS for 10 minutes each, then incubated with secondary antibodies (Jackson Immunoresearch Laboratories) for 1 hour, rinsed 3× with PBS for 10 minutes each, then counterstained with Alexa405-phalloidin (ThermoFisher) for 30 minutes, then rinsed with PBS 3× for 10 minutes each, then mounted with ProLong Glass antifade reagent (ThermoFisher).

### Scanning Electron Microscopy of Membrane Extracted Cells

Cells were briefly washed with 0.1 M phosphate-buffered saline (PBS) (37°C) to remove culture media and immediately treated with a membrane extraction solution containing 1% Triton X-100, 100 mM PIPES (pH 7.2), 1 mM EGTA, 1 mM MgCl2, 4.2 % sucrose, 10 μM taxol (Thermo-Scientific) and 10 μM phalloidin (Sigma) for 10 min with gentle rocking at room temperature. Then the samples were washed twice for 5 min in PBS and fixed with 2% glutaraldehyde, 2% paraformaldehyde (Electron Microscopy Sciences - EMS), in 0.1 M sodium cacodylate buffer for 30 min at RT. Samples were washed in the same buffer, post-fixed with 1% OsO4 (EMS) and 1% tannic acid (EMS) for 1h each, dehydrated with a graded ethanol series until absolute and critical point-dried (Leica CPD 030). The samples were coated with a thin platinum layer (4 nm) (Leica EM SCD500) and imaged on a Zeiss Sigma-VP scanning electron microscope at 5 kV.

### Image processing and analysis

After acquisition, images were Airyscan processed using the auto-filter 2D-SR settings in Zen Blue (ZEISS). All images were post-processed and analyzed using Imaris (BITPLANE) and Fiji software^28^.

### Plasmids

Drp1-mCherry was a kind gift from Gia Voeltz (Addgene plasmid #49152). mCherry-Cyto*b*_5_RR was a gift from Nica Borgese^29^. EGFP-LifeAct was a gift from Jennifer Rohn ^30^. Lamp1-mCherry, SiT-mApple, Sec61-mCherry, and dsRed-Skl were all generous gifts from the Lippincott-Schwartz lab. Rab5a-mCherry was a generous gift from the Merrifield lab (Addgene plasmid #27679). All custom actin nanobody probes were generated starting from the commercial vector of actin chromobody-tagGFP or actin chromobody-tagRFP (ChromoTek) and cloned via the BglII and NotI restriction sites. The following amino acid sequences were attached to the C-terminal of the actin chromobody probes to target the protein either to mitochondria or the ER:

**Fis1 (AC-mito and LifeAct-GFP-Fis1):**

IQKETLKGVVVAGGVLAGAVAVASFFLRNKRR^5^

**Cytb5mito (aka “Cyto*b*_5_RR”) (AC-GFP-Cytb5mito and LifeAct-GFP-Cytb5mito):**

FEPSETLITTVESNSSWWTNWVIPAISALVVALMYRR^31^

**Cytb5ER (AC-ER):**

IDSSSSWWTNWVIPAISAVAVALMYRLYMAED^5^

LifeAct-GFP-Fis1, LifeAct-GFP-Cytb5mito, and AC-GFP-Cytb5mito were generated using PFU Ultra II for megaprimer PCR insertion^32^. The PCR primers, intended modifications, insert templates, and destination plasmids are listed in Supplemental Table 1 below. All constructs were sequenced completely across their coding region.

## Acknowledgements

Many thanks to Andrew Moore (Janelia Farms) and Stephanie Harada (Salk Institute for Biological Studies) for help with the cartoon diagram for Figure 1. We would like to also thank Christopher Obara and Jennifer Lippincott-Schwartz (Janelia Farms) for critical feedback and suggestions on the manuscript. This work was also greatly improved by Twitter feedback on our initial bioRxiv preprint, most especially Subhojit Roy (UW-Madison). The Waitt Advanced Biophotonics Center is funded by the Waitt Foundation and Core Grant applications NCI CCSG (CA014195) and NINDS Neuroscience Center (NS072031). This work was supported by the Transgenic Core Facility of the Salk Institute with funding from NIH-NCI CCSG: P30 014195. R.G. laboratory is funded by grants from HFSP RGP0021/2016 and the Cluster of Excellence CIBSS-Centre for Integrative Biological Signalling Studies. O.A.Q. lab is supported by NIGMS Grant R15 GM119077 and by funding from the University of Richmond School of Arts & Sciences.

---

**Supplemental Table S1:**
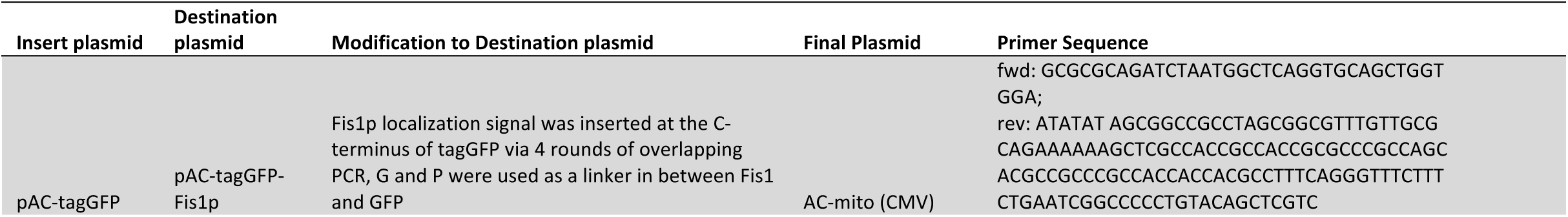
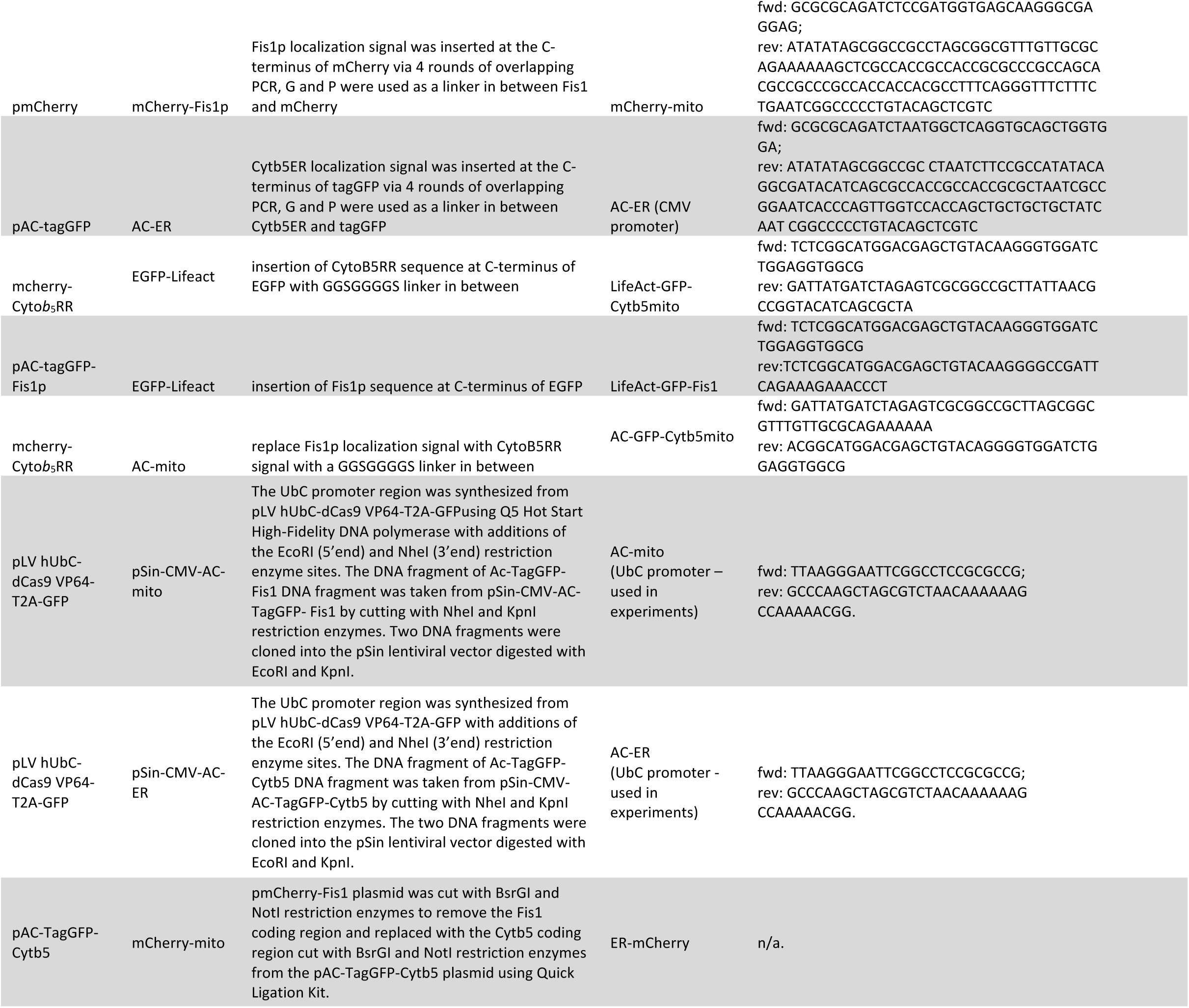
Primers and plasmids used to generate the expression constructs for these studies

**Supplementary Figure 1.**
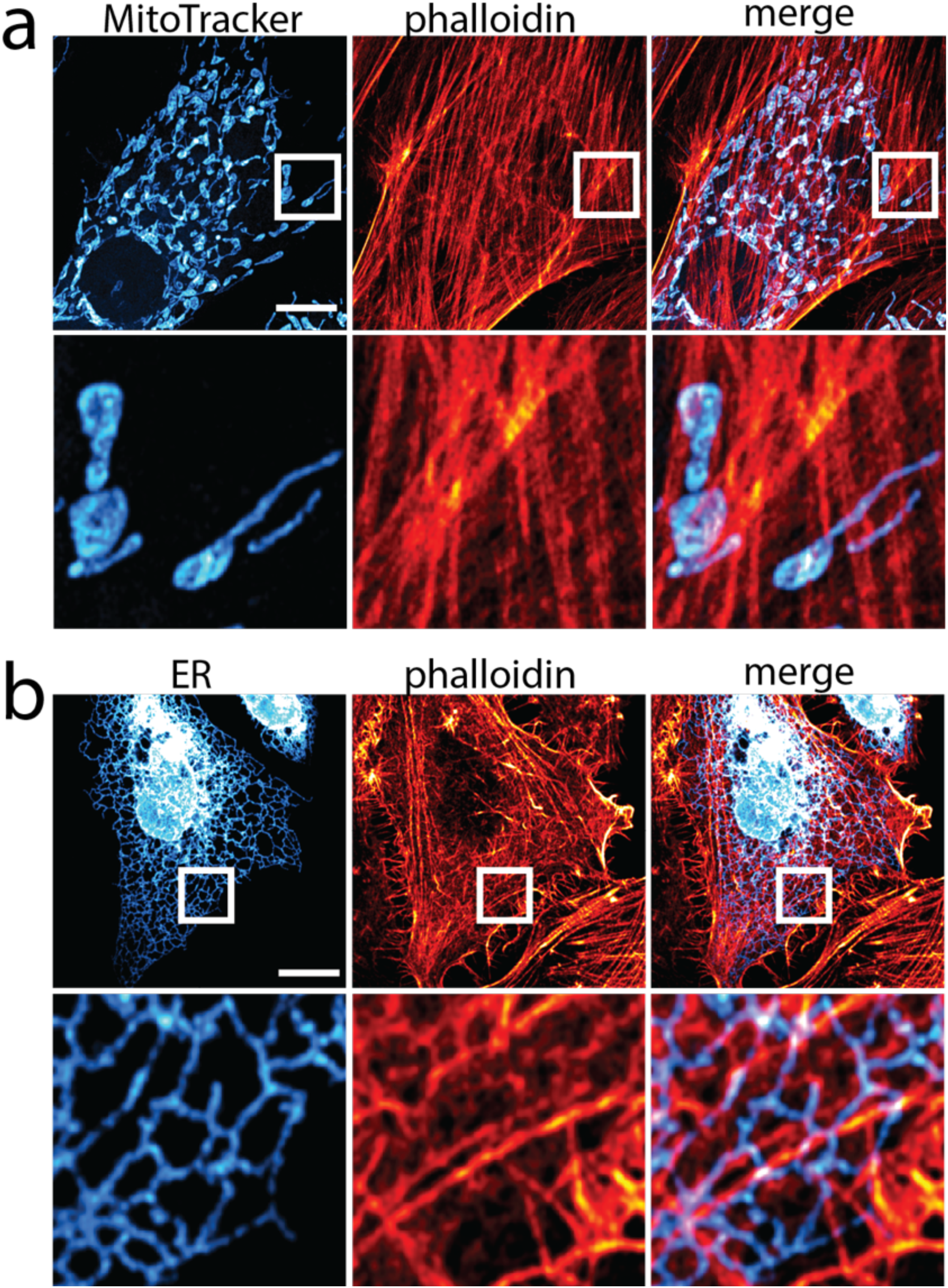
Phalloidin staining does not show obvious accumulation of actin on mitochondria or ER. **a,** Cells stained with phalloidin and MitoTracker do not show obvious accumulation of actin on mitochondria. **b,** Cells expressing the ER marker Sec61-mCherry counterstained with phalloidin do not show obvious accumulation of actin on any specific region of the ER. Scale bar: 10μm.

**Supplementary Figure 2.**
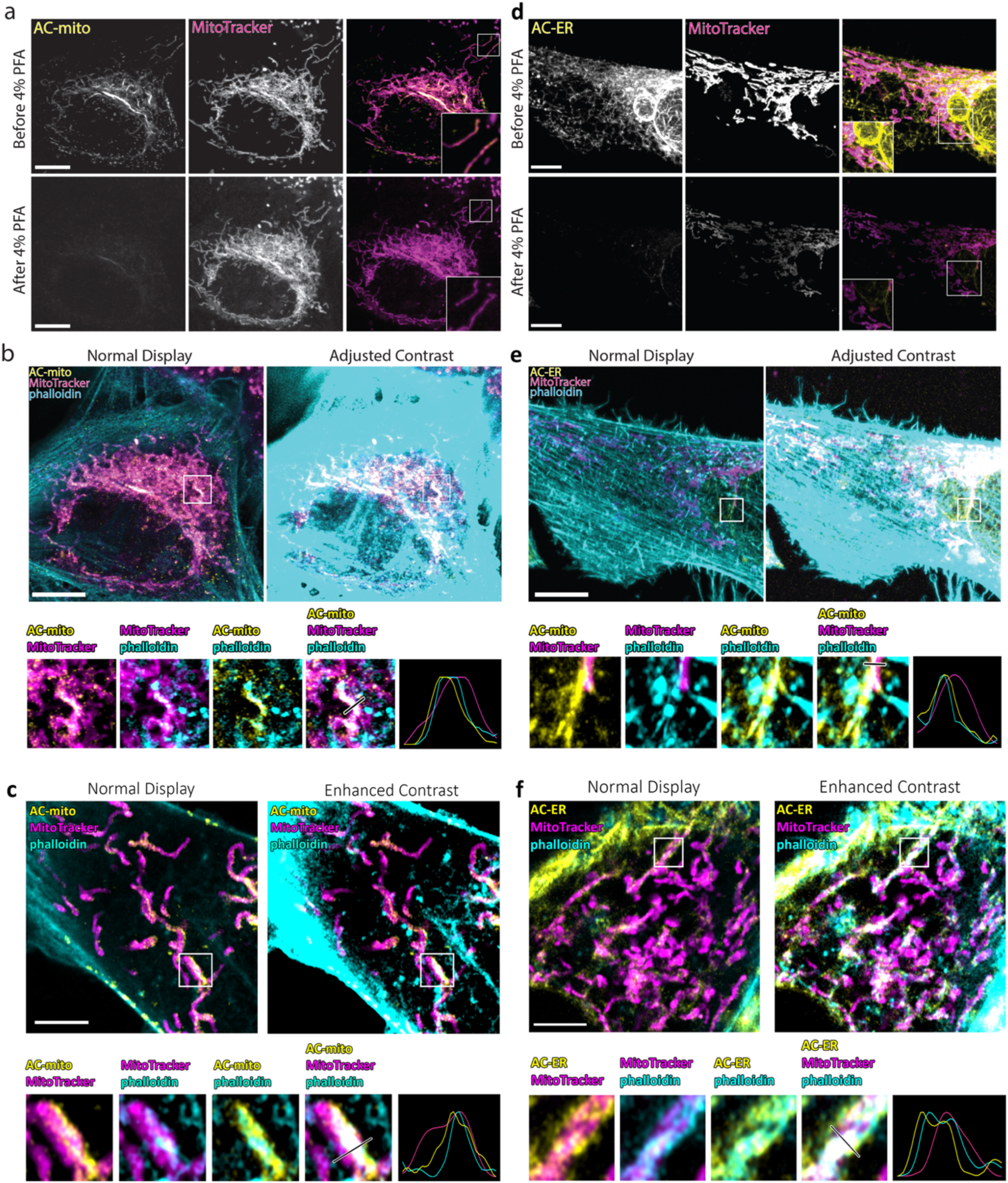
Most sub-organellar actin accumulation does not survive conventional fixation. **a+d** Imaging the same cell expressing AC-mito or AC-ER before and 30 minutes after fixation with 4% PFA reveals a significant decrease in AC-mito or AC-ER signal after fixation. This decrease was observed in all cells imaged (AC-mito: n = 3, AC-ER: n = 4). **b+e,** phalloidin colocalizes with AC-mito and AC-ER, but with much lower signal than surrounding larger actin structures. Bottom row shows insets with adjusted contrast. **c+f,** Cells permeablized with cytoskeleton stabilizing membrane extraction to label phalloidin while preserving AC-mito and AC-ER signal. Phalloidin colocalizes with AC-mito and AC-ER. Scale bars: 10 μm

**Supplementary Figure 3.**
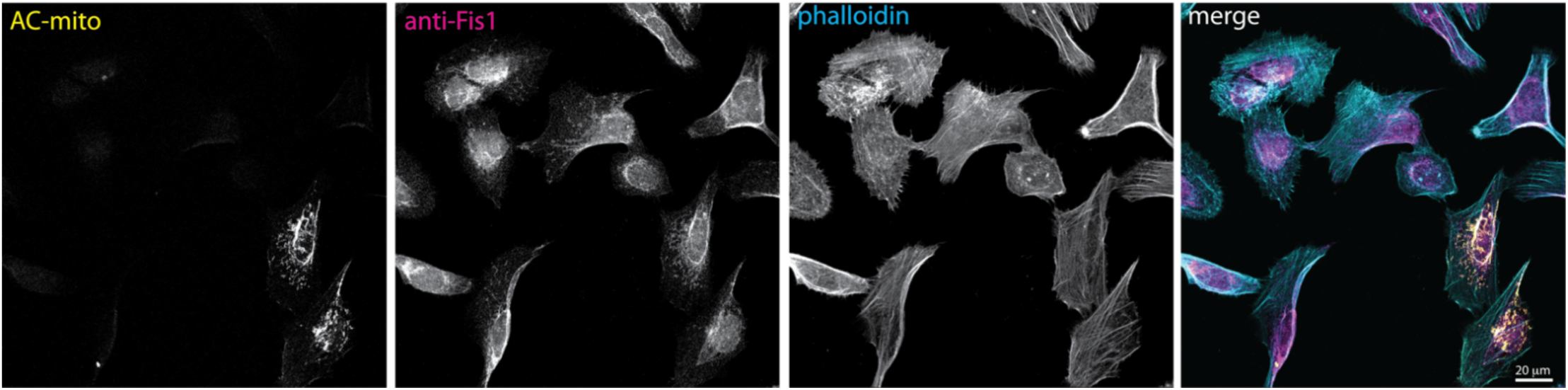
AC-mito expression does not alter endogenous Fis1 localization to mitochondria. Cells expressing AC-mito show similar levels of endogenous anti-Fis1 immunofluorescence as non-transfected cells. Note that the anti-Fis1 antigen is not present in the AC-mito protein. Scale bar: 20 μm.

**Supplementary Figure 4.**
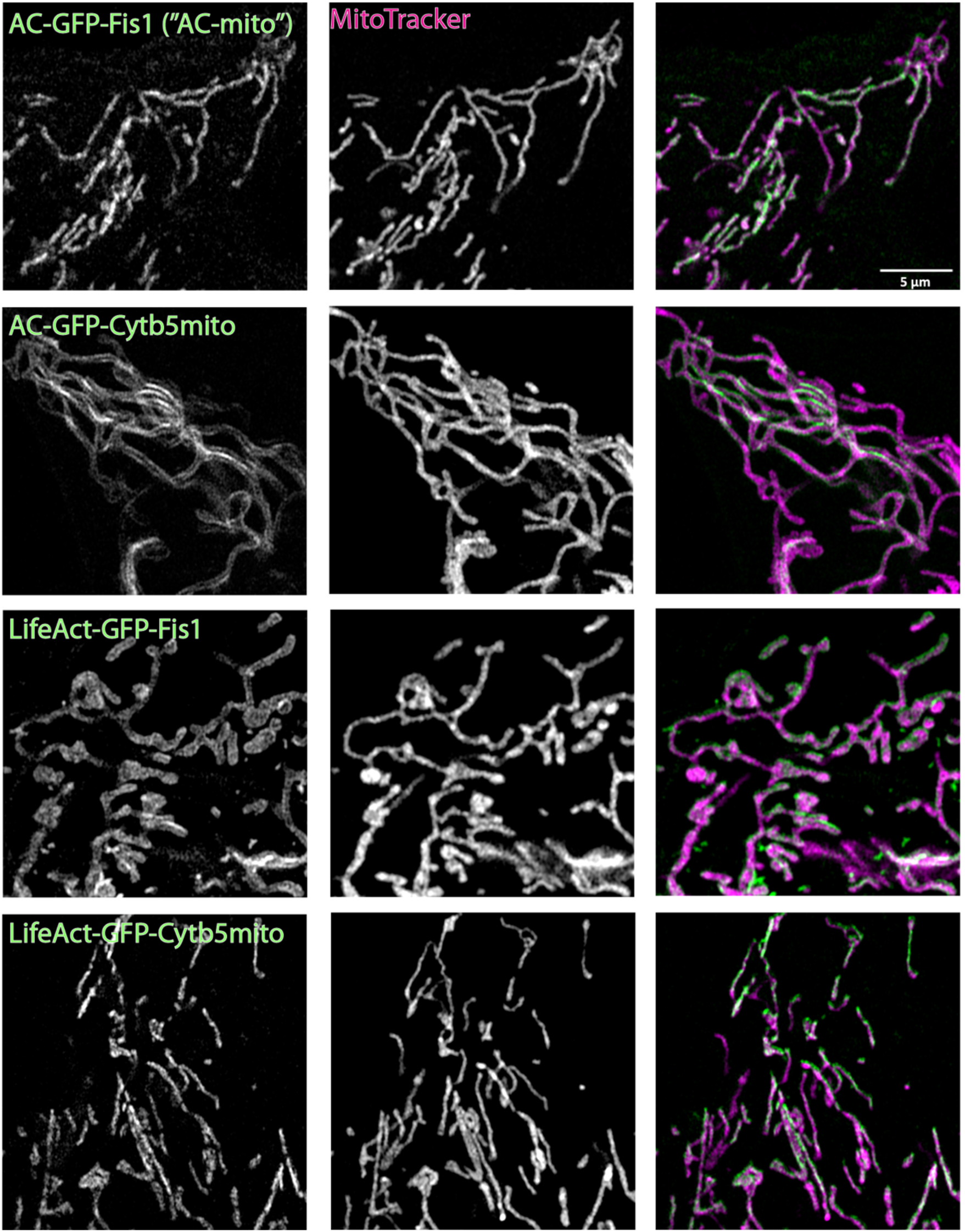
Alternate membrane-targeting and actin-binding motifs yield similar results. Switching the Fis1 mitochondrial outer membrane targeting sequence for Cytb5mito (Cytb5mito-AC-GFP) yields similar results. Similarly, switching the actin nanobody motif with LifeAct (Fis1-LifeAct-GFP) also yields similar results. Scale bar: 5 μm.

**Supplementary Figure 5.**
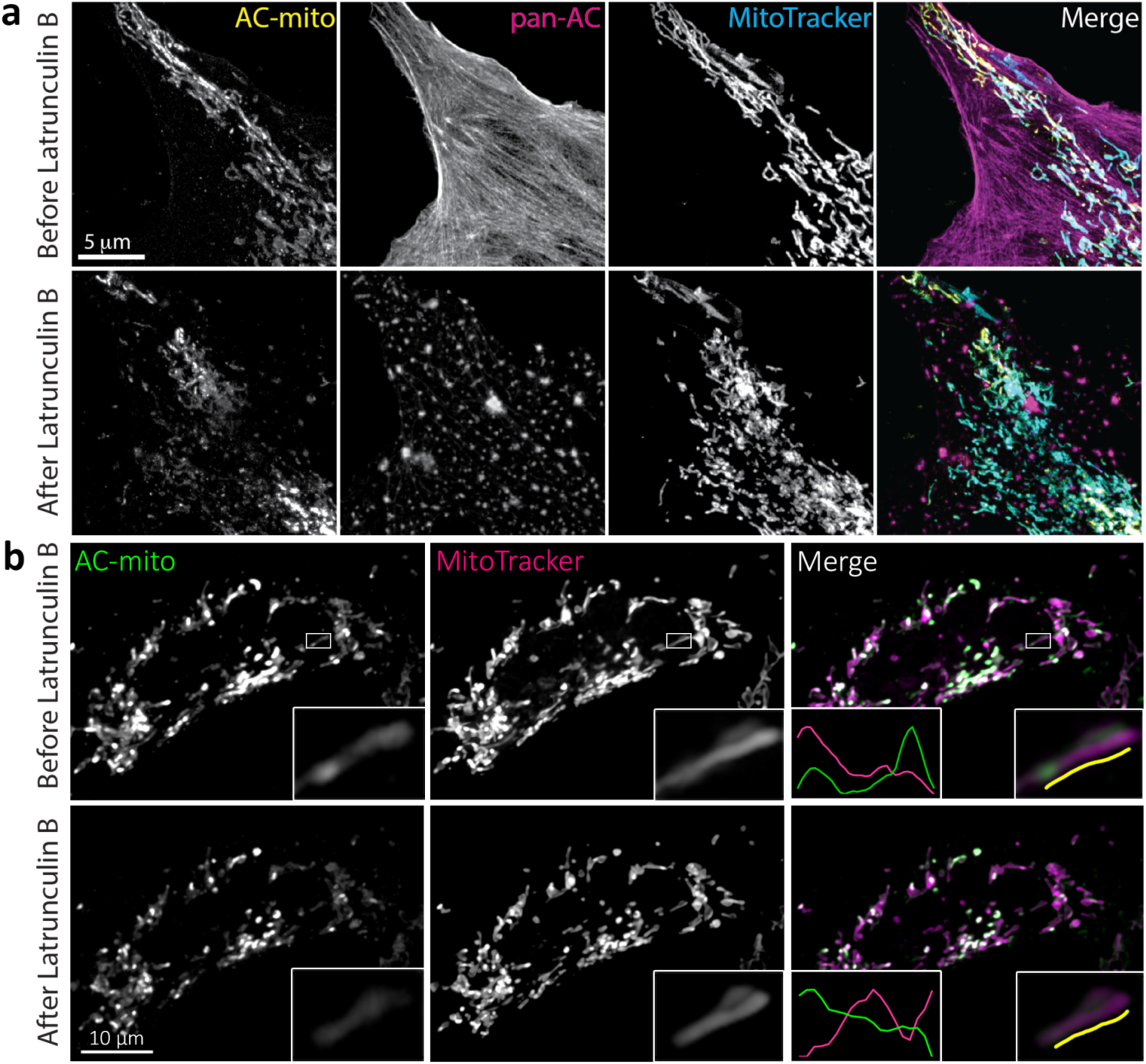
Actin depolymerization destroys AC-mito localization. **a,**Cells expressing AC-mito and pan-AC (ACtagRFP) were imaged before and after treatment with Latrunculin B for 30 minutes. After Latrunculin B treatment, AC-mito displayed much less accumulation at specific regions on mitochondria. This loss of accumulation was observed in all cells imaged after treatment with Latrunculin B or Cytochalasin D (n = 6). Scale bar: 5 μm **b,** Cells expressing AC-mito before and after treatment with Latrunculin B including example insets. The more diffuse localization of AC-mito following Latrunculin B treatment is apparent in the example mitochondrion. Scale bar: 10 μm

**Supplementary Figure 6.**
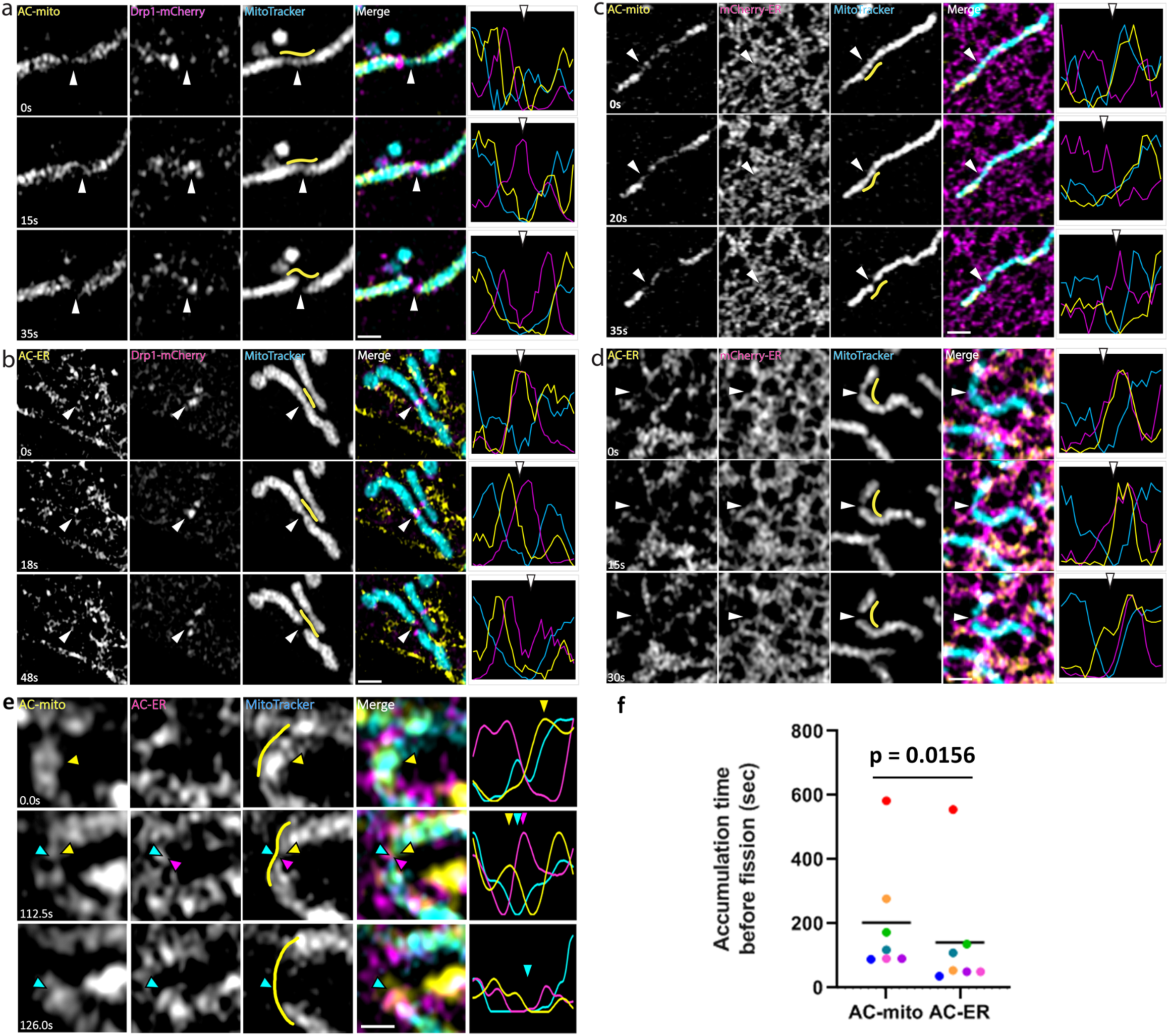
Live imaging of AC probes reveal actin accumulation at mitochondrial fission sites. **a,** AC-mito accumulates prior to Drp1 during mitochondrial fission. **b,** AC-ER accumulates at mitochondrial fission sites with Drp1. **c,** AC-mito accumulates at ER-mitochondria contact sites prior to mitochondrial fission. **d,** AC-ER specifically accumulates at ER-mitochondria contact sites prior to mitochondrial fission. **e,** AC-mito accumulates at mitochondrial fission sites prior to AC-ER accumulation. Scale bars: 1μm. **f,** Quantification of e. p-value determined by paired ratio t-test.

**Supplementary Figure 7.**
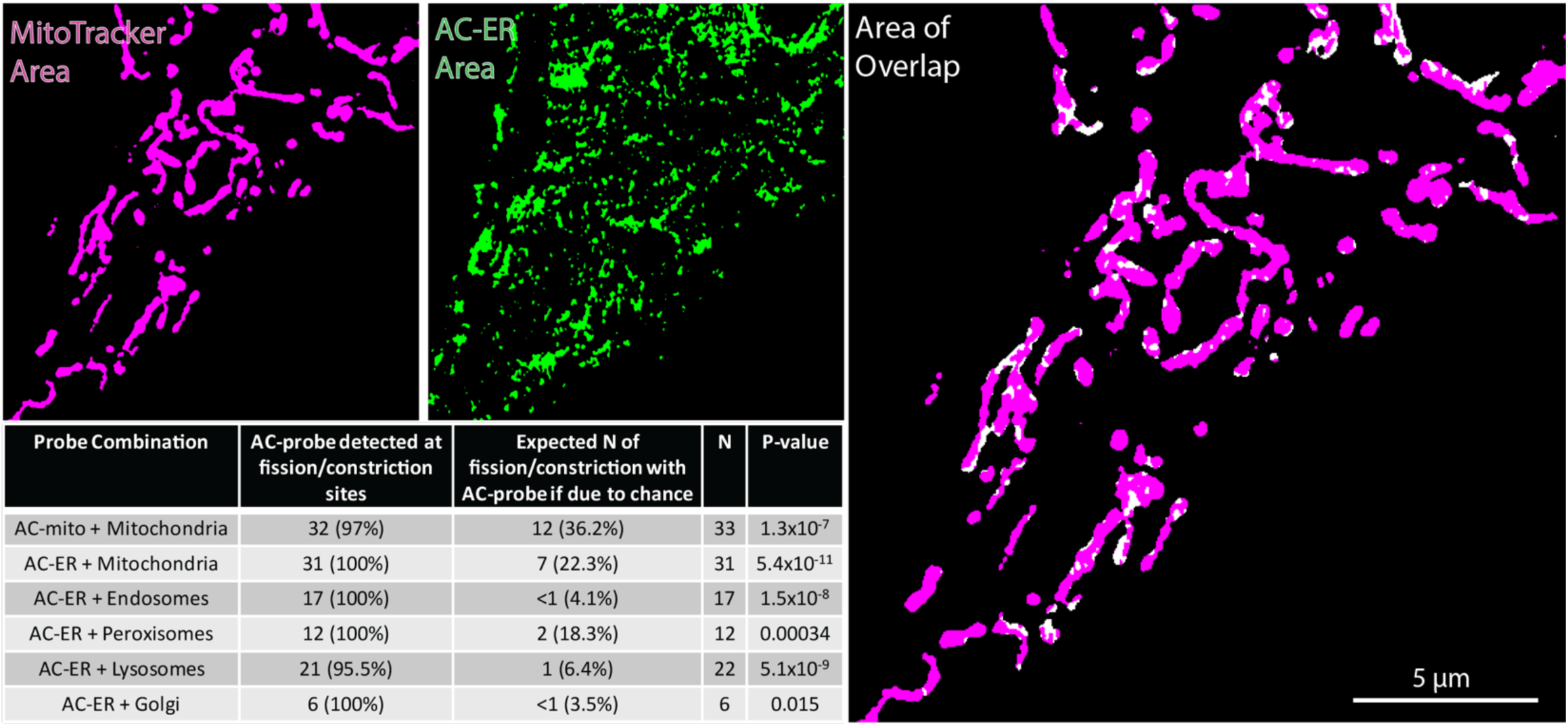
AC-mito and AC-ER accumulate at sites of organelle fission/constriction at a significantly higher frequency than would be expected by chance. Images show an example of the masked areas of MitoTracker, AC-ER, and areas of overlap. The table displays the observed frequency of AC probes observed at fission/constriction sites compared to the frequency that would be expected by chance. P-values determined by Fisher’s exact test.

**Supplementary Movie 1|Live cell imaging of mitochondria-associated actin.** A Hap1 cell expressing AC-mito counterstained with MitoTracker shows dynamic subdomains of actin enrichment on mitochondria.

**Supplementary Movie 2|Live cell imaging of mitochondria-associated actin.** A Hap1 cell expressing AC-mito counterstained with MitoTracker shows dynamic subdomains of actin enrichment on mitochondria (example 2).

**Supplementary Movie 3|Live cell imaging of ER-associated actin.** A Hap1 cell expressing AC-ER counterstained with MitoTracker shows dynamic subdomains of actin enrichment on the ER, in particular at ER-mitochondria contact sites.

**Supplementary Movie 4|Live cell imaging of ER-associated actin.** A Hap1 cell expressing AC-ER counterstained with MitoTracker shows dynamic subdomains of actin enrichment on the ER, in particular at ER-mitochondria contact sites (example 2).

**Supplementary Movie 5|FRAP of cell co-expressing AC-mito and mCherry-mito.** FRAP of U2OS cells co-expressing AC-mito and mCherry-mito shows that AC-mito is highly mobile but less mobile than mCherry-mito.

**Supplementary Movie 6|FRAP of cell co-expressing AC-ER and mCherry-ER.** FRAP of U2OS cells co-expressing AC-ER and mCherry-ER shows that AC-ER is highly mobile but less mobile than mCherry-ER.

**Supplementary Movie 7|Live imaging of AC-mito expressing cells before and after Cytochalasin D treatment.** Live imaging of Cytochalasin D untreated (left) vs. treated (right) AC-mito expressing U2OS cells.

**Supplementary Movie 8|Live imaging of AC-mito and Drp1 during mitochondrial fission.** Live imaging of AC-mito and Drp1-mCherry in U2OS cells counterstained with MitoTracker reveals accumulation of mitochondria-associated actin prior to Drp1-mediated fission.

**Supplementary Movie 9|Live imaging of AC-ER and Drp1 during mitochondrial fission.** Live imaging of AC-ER and Drp1-mCherry in U2OS cells counterstained with MitoTracker reveals accumulation of ER-associated actin prior to Drp1-mediated fission. Scale bar: 1 μm.

**Supplementary Movie 10|Live imaging of AC-mito and the ER during mitochondrial fission.** Live imaging of AC-mito and mCherry-ER in U2OS cells counterstained with MitoTracker reveals accumulation of mitochondria-associated actin prior to ER-mediated fission. Scale bar: 1 μm.

**Supplementary Movie 11|Live imaging of AC-ER and the ER during mitochondrial fission.** Live imaging of AC-ER and mCherry-ER in U2OS cells counterstained with MitoTracker reveals accumulation of ER-associated actin prior to ER-mediated fission. Scale bar: 1 μm.

**Supplementary Movie 12|Live imaging of AC-mito and AC-ER during mitochondrial fission.** Live imaging of AC-mito and AC-ER in U2OS cells counterstained with MitoTracker reveals accumulation of mitochondria-associated actin prior to accumulation of ER-associated actin. Scale bar: 1 μm.

**Supplementary Movie 13|ER-associated actin accumulates at mitochondrial fission sites.** AC-ER accumulation at a fission site in MitoTracker-labeled mitochondria. Scale bar: 1 μm

**Supplementary Movie 14|ER-associated actin accumulates at endosomal fission sites.** AC-ER accumulation at two fission sites in Rab5a-mCherry-labeled endosomes. Scale bar: 1 μm

**Supplementary Movie 15|ER-associated actin accumulates at peroxisomal fission sites.** AC-ER accumulation at a fission site in dsRed-Skl-labeled peroxisomes. Scale bar: 1 μm

**Supplementary Movie 16|ER-associated actin accumulates at lysosomal fission sites.** AC-ER accumulation at a fission site in LAMP1-mCherry-labeled lysosomes. Scale bar: 1 μm

**Supplementary Movie 17|ER-associated actin accumulates at Golgi fission sites.** AC-ER accumulation at a fission site in SiT-mApple-labeled Golgi. Scale bar: 1 μm

**Supplementary Movie 18|ER-associated actin forms a network on the nucleus.** 3D rendering of a U2OS cell expressing AC-ER reveals accumulation around the surface of the nucleus. Cyan: AC-ER. Orange: MitoTracker.

